# Demographic feedbacks can hamper the spatial spread of a gene drive

**DOI:** 10.1101/2021.12.01.470771

**Authors:** Léo Girardin, Florence Débarre

## Abstract

This paper is concerned with a reaction–diffusion system modeling the fixation and the invasion in a population of a gene drive (an allele biasing inheritance, increasing its own transmission to offspring). In our model, the gene drive has a negative effect on the fitness of individuals carrying it, and is therefore susceptible of decreasing the total carrying capacity of the population locally in space. This tends to generate an opposing demographic advection that the gene drive has to overcome in order to invade. While previous reaction–diffusion models neglected this aspect, here we focus on it and try to predict the sign of the traveling wave speed. It turns out to be an analytical challenge, only partial results being within reach, and we complete our theoretical analysis by numerical simulations. Our results indicate that taking into account the interplay between population dynamics and population genetics might actually be crucial, as it can effectively reverse the direction of the invasion and lead to failure. Our findings can be extended to other bistable systems, such as the spread of cytoplasmic incompatibilities caused by Wolbachia.

## 1. Introduction

### 1.1. Biological background

Some genetic elements can bias inheritance in their favor, and therefore spread in a population over successive generations, even if they are costly to the organism carrying them [40], a phenomenon called gene drive [1]. Inspired from natural systems, synthetic gene drives have been developed over the last decades for the control of natural populations [33]. Artificial gene drives can be used to spread a genetic modification in a natural population [8], in order to *(i)* modify this target population (for instance, make disease vectors resistant to the pathogen that they transmit) without significantly affecting its size (replacement drives), or *(ii)* reduce the size of the target population by spreading an allele that affects fecundity or survival (suppression drives), or even eradicate the target population (eradication drives) [12].

One way of biasing transmission involves copying the drive element onto the other chromosome in heterozygous cells [7], a process called gene conversion or “homing”. A target sequence on the other chromosome is recognized, cleaved, and a DNA-repair mechanism using the intact chromosome as a template leads to the duplication of the drive element. The drive element is then present in most gametes and therefore in most offspring. The high flexibility and programmability brought about by the CRISPR-Cas genome editing technology greatly simplifies the creation of this type of artificial gene drives [16], and has led to a renewed interest in this technique of population control.

To date, artificial gene drives have been confined to laboratory experiments; field trials, whether small-scale and confined, or open, have not taken place yet [15]. In between, mathematical models can help identify features of gene drives that are most important in determining their dynamics; models can also help anticipate potential issues, and are an essential step between lab and field experiments [27].

A key feature affecting the spread of gene drives is the existence or not of a release threshold. Some drive systems spread from arbitrary low introduction frequencies (“threshold-independent drives”), while some others only do if enough drive-carrying individuals are introduced in the population (“threshold-dependent drives” or “high-threshold drives”) [15, p19]. In mathematical terms, a release threshold corresponds to the existence of an unstable internal equilibrium (*i*.*e*., where wild-type and drive alleles coexist), in other words, of a bistability. Homing-based gene drives can belong to the two categories, depending on their intrinsic parameters [11].

The existence of a release threshold affects the spatial spread of a drive and the potential for spatial confinement. High-threshold drives can be spatially confined if dispersal is limited [2, 21, 24]. In spatially continuous environment, the shape of the wave of advance of a drive system changes from pulled for no-threshold drives to pushed in high-threshold drives [12]. A wave of advance of a high-threshold drive can be stopped by obstacles [36]. More generally, these results are in line with those of models of spatial spread of bistable systems [3, 4].

Many mathematical models of the spatial spread of gene drives focus on allele frequencies and ignore changes in population density. Changes in the size of the target population will however happen with the spread of a gene drive, either as a potential side effect for replacement drives, or as a feature for suppression and eradication drives. Considering these spatial and temporal variations in population size matters, because they create fluxes of individuals from more densely to less densely populated locations, opposing the spread of the gene drive [12].

Beaghton *et al*. [5] used a reaction–diffusion framework to study the spatial spread of a driving-Y chromosome causing population suppression or eradication, and explicitly followed population densities. In a driving-Y system, the offspring of driving-Y bearing males are almost all driving-Y bearing males, because the development of X-chromosome bearing gametes is disrupted. A driving-Y is a no-threshold drive: in a well-mixed population, a driving-Y system can increase from arbitrary low frequencies. Beaghton *et al*. showed that a driving-Y system would spread spatially, and they calculated the speed of the wave of advance.

In this article, we use a system of partial differential equations to study the spread of a homing-based gene drive over a one-dimensional space, explicitly taking into account changes in population sizes. In our model, the fitness cost *s* associated to the drive affects the level of population suppression, but also the existence and value of the release threshold. We explore the model numerically and prove some results mathematically for large regions of the parameter space. We explore the robustness of our finding by considering other forms of density dependence (including Allee effects), assuming fitness effects on other fitness components, and finally extend our result to another bistable system, the spread of cytoplasmic incompatibilities brought about by *Wolbachia*.

### 1.2. Organization of the paper

In the rest of Section 1, we present our mathematical model, relate it to the literature, and state our results. In Section 2, we explain our numerical method and prove our analytical results. In Section 3, we conclude with a mathematical and biological discussion.

### 1.3. Derivation of the reaction–diffusion system

To derive our equations, we follow the methodology presented in [31]. In addition, we assume that heterozygous individuals are functionally equivalent to drive homozygotes. This can be the case when the conversion rate is 100%, *and* either *(i)* gene conversion (also called homing) takes place early in development (typically in the zygote – as opposed to gene conversion in the germline) – so that heterozygotes are all immediately converted into homozygotes, and no heterozygous individuals are ever introduced to the population from an outside source, or *(ii)* gene conversion takes place later in the life cycle, but the drive is dominant, so that heterozygotes have the same fitness as drive homozygotes (and then only transmit the drive allele, since gene conversion is 100%). This assumption simplifies a lot the model since we have to track two genotypes instead of three in general: in scenario *i*), there are never heterozygous individuals in the population, and in scenarion *ii*), heterozygotes can be conflated with drive homozygotes since they behave exactly the same way. This leads to the following reaction–diffusion system:

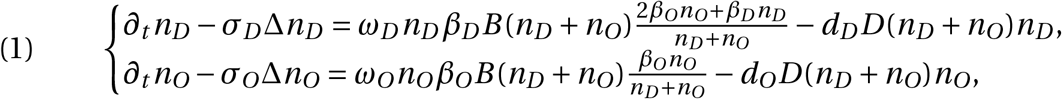

where:

- *n*_*D*_ (*t, x*) and *n*_*O*_(*t, x*), nonnegative functions of time *t* and space *x*, are (continuous densities approximating) the number of alleles *D* and *O* respectively, or (up to a factor 1/2) the number of homozygous individuals *DD* and *OO* respectively;
- for each allele *a* ∈ {*D,O*}, *σ*_*a*_ is the spatial diffusion rate of homozygous *aa* individuals, *ω*_*a*_ is their juvenile survival rate (the proportion of newborns that survive until the adult age), *β*_*a*_ is the fecundity rate of a gamete carrying *a* (fecundity being defined here multiplicatively in the sense that a mating between *aa* and *AA* individuals will produce *β*_*a*_*β*_*A*_ newborns), *d*_*a*_ is the death rate of *aa* individuals, and all these parameters are positive constants;
- *B* and *D*, nonnegative functions of the total population size, are the per capita birth rate and death rate. If they were positive constants, then a pure wild-type population would undergo exponential growth, which is not realistic; by using non-constant, *a priori* nonlinear, functions, we can model more complex population dynamics, like logistic growth or Allee effects.

Our assumptions of early and 100% successful gene conversion (homing) are reflected in the presence of the +2*β*_*O*_*n*_*O*_ term in the first equation of system (1); assuming late gene conversion and a dominant drive allele, together with 100% gene conversion, results in the same equations.

Defining

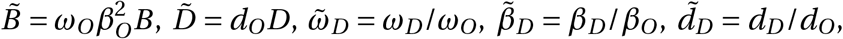

we obtain the reduced system:

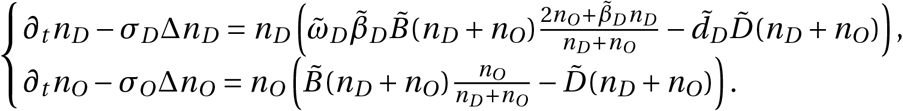

Assuming subsequently that selection does not act on mobility (*σ*_*D*_ *= σ*_*O*_), changing the variable *x* into 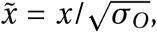 and getting rid of all “∼” for ease of reading, we obtain the reduced system:

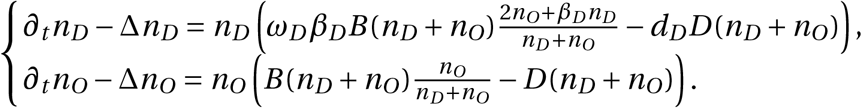

The biological meaning of the parameters in this simpler system is the following: *ω*_*D*_, *β*_*D*_, *d*_*D*_ are multiplicative variations to the “norm” fixed by *OO* individuals. For instance, if *β*_*D*_ = 1/2, then wild-type homozygotes (*OO*) are twice more fecund than drive homozygotes (*DD*).

Finally, when selection only acts on survival with a *fitness cost s* ∈ [0, 1] (*d*_*D*_ *= β*_*D*_ = 1, *ω*_*D*_ = 1 − *s*, standing assumptions from now on), the system reads:

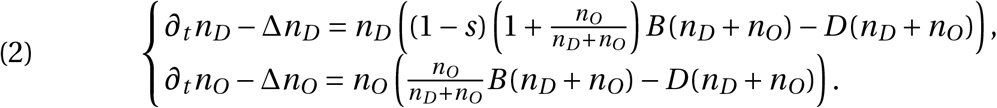

Defining *n*(*t, x*) *= n*_*D*_ (*t, x*) + *n*_*O*_(*t, x*) the total population density and 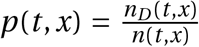 the proportion of allele *D*, we find that the equivalent system satisfied by (*p, n*) is:

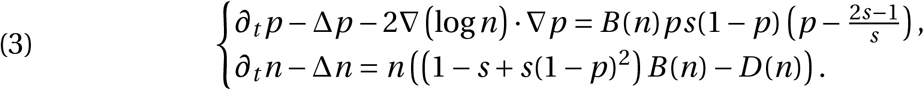

In the above system and in the whole paper, log is the natural logarithm.

*B*(*n*) and *D*(*n*) are still unspecified at this point.

The equation for *p* in system (3) has a non-trivial equilibrium 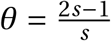, which is admissible when 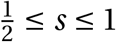. In this case, this internal equilibrium is unstable. The release-threshold properties of the gene drive are therefore given by the value of *s*: the drive is threshold-independent when 0 ≤ *s* ≤ 1/2, and threshold dependent when 1/2 < *s* ≤ 1, the value of the release threshold being *θ*.

### 1.4. Carrying capacity and gene drive typology

A pure wild-type population is governed by the equation *∂*_*t*_ *n*_*O*_−Δ*n*_*O*_ *= n*_*O*_(*B*(*n*_*O*_)−*D*(*n*_*O*_)). Hence *B*−*D* can be understood as the wild-type intrinsic growth rate per capita. Subsequently, we define the *wild-type carrying capacity* (*i*.*e*., equilibrium population size) as follows:

- if *B* (*n*)−*D*(*n*) < 0 for any *n* ≥ 0, then the carrying capacity is 0;
- otherwise, the carrying capacity is the maximal value of *n* ≥ 0 for which *B* (*n*) − *D*(*n*) ≥ 0.

For well-posedness purposes, we assume that the carrying capacity is finite, and without loss of generality we assume that it is 1, fixing the unit of population density^1^.

In a spatially homogeneous setting, a population goes extinct if, and only if, its carrying capacity is zero. The growth rate of a pure *DD* population is (1−*s*)*B* (*n*_*D*_)−*D*(*n*_*D*_). Hence, in our framework,

- *s =* 0 corresponds to pure, costless, replacement drives;
- as soon as *s* > 0, the drive is a suppression drive, lowering the carrying capacity of the population;
- the necessary and sufficient condition for the drive to be an eradication drive, namely a drive whose carrying capacity is 0, is:

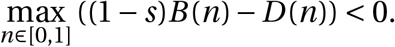

Finally, a simple rescaling of space and time shows that the pair (*B, D*) and the pair (*αB, αD*), for any *α* > 0, lead to the same system. In order to fix the ideas, in what follows, the pair (*B, D*) is normalized by the standing assumption min_*n*∈[0,1]_ *D*(*n*) = 1.

### 1.5. Relations with other models from the literature

In the recent literature, reaction– diffusion models related to system (4) have attracted a special attention, be it for the study of gene drive or in other evolution or population genetics contexts ^2^. Let us show how our model relates to some of those earlier models.

Fixing in the following discussion *D*(*n*) = 1, system (4) becomes: (4)

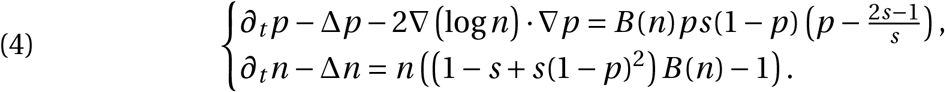

#### 1.5.1. Deriving ref. [20, 26, 36] from system (4)

Recall that in this discussion, *D*(*n*) = 1.

If *B* (*n*) is a positive constant (population subjected to pure, unrealistic, Malthusian growth), then up to a rescaling of (*t, x*), the equation on *p* reads

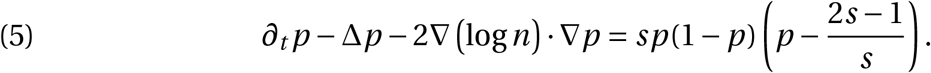

Together with V. Calvez, we studied this equation in a gene drive context [20, Section 4.3] without *a priori* knowledge on *n*. It has also been studied in a mosquito–Wolbachia context by Nadin, Strugarek and Vauchelet in [26].

Back to (4) with a non-constant function *B*, recalling that the wild-type carrying capacity is 1 (namely, *B* (1) = 1), we consider small variations about *n =* 1 (a replacement drive) by defining a “rescaled total population” 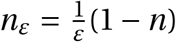 (*i.e., n* = 1 − *εn*_*ε*_) and, following Strugarek–Vauchelet [35], we find

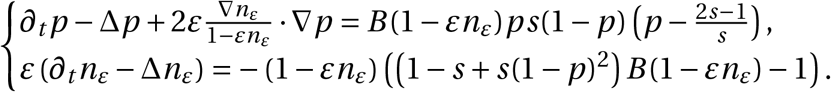

Formally, if *ε* → 0 with the scalings 1− *εn*_*ε*_ ↛ 0, *ε∂*_*t*_ *n*_*ε*_ → 0, *ε*∇*n*_*ε*_ → 0, *ε*Δ*n*_*ε*_ → 0, this system becomes

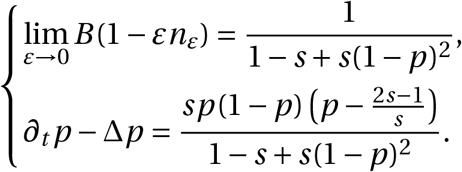

This limit can actually be made rigorous [35] provided:

- the birth rate *B = B*_*ε*_ depends on the small parameter *ε* in such a way that *B*_*ε*_(1− *εn*_*ε*_) can be rewritten as a function of *n*_*ε*_ only, *i*.*e*.,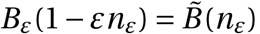;
- the time horizon is finite, or in other words, we are only concerned with the early gene drive dynamics, and not with its long-time asymptotic spreading.

It turns out that the latter equation on *p* is exactly the equation studied in a gene drive spreading context by Tanaka–Stone–Nelson [36]:

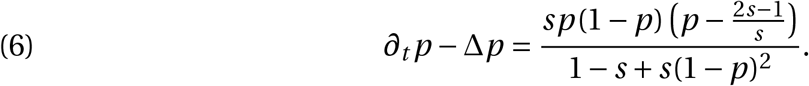

Although Tanaka–Stone–Nelson’s paper [36] and Strugarek–Vauchelet’s paper [35] were published the same year and without direct knowledge one from another, they can therefore be related *a posteriori*: our general model reduces to Tanaka–Stone–Nelson’s model via Strugarek–Vauchelet’s limiting procedure in a context of replacement drives invading populations at carrying capacity, and Tanaka–Stone–Nelson’s model can be used to study spreading properties provided the Strugarek–Vauchelet limiting procedure remains valid with an infinite time horizon. In this regard, our paper can be understood as an investigation of the validity of the approximation (6) to study spreading properties and for suppression or eradication drives.

Back to system (4), it is formally clear that, under a weak selection assumption *s* ≃ 0, the total population *n* does not depend a lot on the allelic frequency *p*, is close to the wild-type carrying capacity 1 everywhere, and by virtue of *B*(1) = 1, the equation on *p* can be approximated by

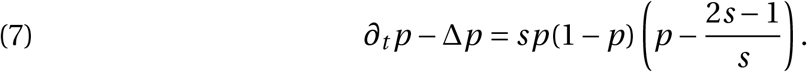

This equation is both an approximation of (5) with *n* ≃ 1 and an approximation of (6) with *s* ≃ 0. This formal reasoning illustrates in another way how our model is consistent with the model of Tanaka–Stone–Nelson [36] under a weak selection assumption ^3^.

Finally, another variant might be (8)

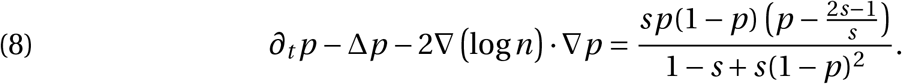

The difference with eq. (6) is the presence of the advection term 2∇(log *n)* · ∇*p*. In our paper with V. Calvez [20], we also studied equation (8), without *a priori* knowledge on *n*. It is not clear how this equation can be deduced from our current model and we leave this as an open problem – yet, in the next section we will explain how all these models descend from a unique discrete-time model. In any case, our intent there was to depart from the Tanaka–Stone–Nelson model and to introduce basic population density effects by means of this advection term. As such, this first work motivated the present one and this modeling clarification as well.

#### 1.5.2. Deriving all models from a unique discrete-time one

Alternatively, we can derive all these models from a discrete-time model with non-overlapping generations [11, 38, 39].

To obtain (4), the operations have to be performed in the following order:

1. write the equations for *n*_*D*_ (*t* + 1) and *n*_*O*_(*t* + 1);
2. subtract from it *n*_*D*_ (*t*) and *n*_*O*_(*t*) respectively and perform a first-order Taylor expansion *n*_*D,O*_(*t* + 1) − *n*_*D,O*_(*t*) ≃ *∂*_*t*_ *n*_*D,O*_;
3. add spatial diffusion to the equations;
4. then write the equivalent system satisfied by (*p, n*); it is (4) indeed.

Going back to refs. [20, 36], we find that there the operations are performed in the following different order:

1. write the equations for *n*_*D*_ (*t* +1) and *n*_*O*_(*t* +1);
2. write the equivalent system satisfied by (*p*(*t*+1), *n*(*t*+1));
3. by Taylor expansion, write the continuous-time system satisfied by (*p, n*);
4. add spatial diffusion (with [20] or without [36] the gene flow advection term).

Although the obtained equation on *n* remains the same as in (4), the equation on *p* differs and is (6) or (8) (depending on whether the advection term was accounted for).

Let us highlight the fact that the two processes above lead to different systems and are not equivalent. Nevertheless, we pointed out in [20, Section 4.3] that the two systems have similar qualitative behaviors.

### 1.6. Main results

In what follows, we choose to focus, for the sake of exposition, on the simplest non-constant choice for *B* and *D*, namely a constant death rate with logistic wild-type growth: *D*(*n*) = 1, *B* (*n*) *= r* (1 − *n*) + 1. The parameter *r* >0 is the intrinsic growth rate of a wild-type population, or *wild-type intrinsic growth rate* for short (in the literature, intrinsic growth rates are also sometimes known as Malthusian growth rates). Thus we have two parameters in the model, the fitness cost *s* and the wild-type intrinsic growth rate *r*. We will discuss more complicated choices in Section 3.

The reaction–diffusion systems are:

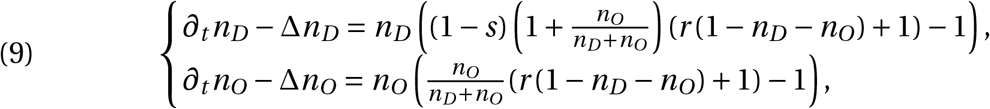

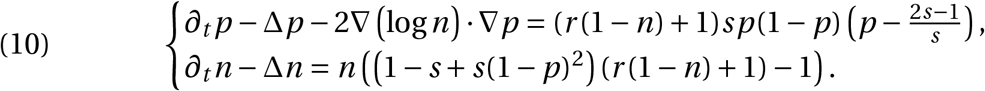

For these systems, the eradication condition is 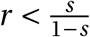.

Since we are concerned with spreading properties, and more precisely with the ability of the gene drive to invade a wild-type population when introduced in sufficiently high numbers in a confined area of space, we neglect boundary effects, assume that the physical space is a Euclidean space – to simplify even further, we assume it is the one-dimensional Euclidean space ℝ – and assume that the initial conditions are close to *n*(0, *x*) = 1 everywhere and *p*(0, *x*) = 1 in a sufficiently large compact interval.

#### 1.6.1. Numerical results

All numerical simulations are performed in *GNU Octave* [14] using standard implicit finite difference schemes. Two examples of numerical code, that can be used to obtain Figure 1 and Figure 4c, are presented online at https://plmlab.math.cnrs.fr/fdebarre/2021_DriveInSpace.

**Figure 1.**
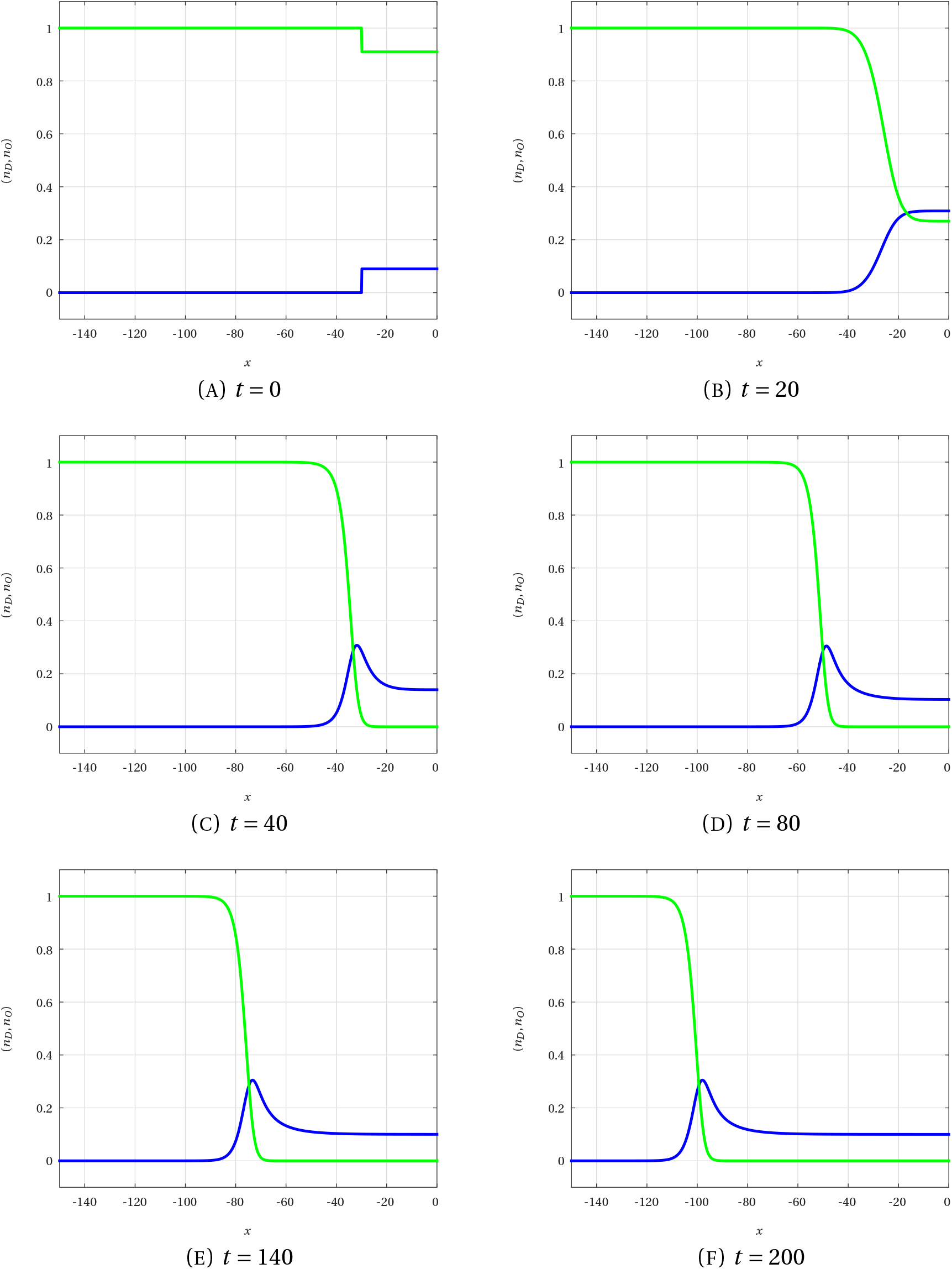
Numerical simulation of the solution of the system (9) at different times (varying time between two snapshots). Blue curve: *n*_*D*_ (*t, x*). Green curve: *n*_*O*_(*t, x*). Here *r =* 10/9, *s =* 0.5, so that the drive is not an eradication drive (1 − *s*/(*r* (1 − *s*)) = 0.1) and it is threshold-independent ((2*s* − 1)/*s =* 0).

To begin with, we simulate the evolution in time of the population densities *n*_*O*_(*t, x*) and *n*_*D*_ (*t, x*).

The spatial domain ℝ is, as usual, approximated by a very large bounded domain with Neumann boundary conditions. Our initial condition is such that (*n*_*D*_, *n*_*O*_) = (0, 1) on the left and (*n*_*D*_, *n*_*O*_) = (0.1, 0.9) on the right.

Time snapshots of such a simulation with *r =* 10/9 and *s =* 1/2 are presented in Figure 1. We clearly observe the rapid convergence of the solution to a *traveling wave*, namely to a solution (*n*_*D*_, *n*_*O*_)(*t, x*) = (*N*_*D*_, *N*_*O*_)(*x* −*ct*) with constant profile and constant wave speed *c* < 0. From now on, we use capital letters *N*_*D*_, *N*_*O*_, *N* and *P* for the wave profiles, functions of *x* −*ct*, and lower-case letters *n*_*D*_, *n*_*O*_, *n* and *p* for the population densities, functions of (*t, x*).

Note that by symmetry and isotropy, we would obtain an axis-symmetric figure if the gene drive was introduced on the left. If it was introduced in the center of a twice as large interval, we would observe the gluing of two traveling waves, one spreading towards the left at speed *c* < 0 and the other spreading towards the right at speed − *c* > 0.

With this choice of parameters *r* and *s*, we observe the so-called *hair-trigger effect*, namely the fact that no matter how small the height and the width of the initial *n*_*D*_, invasion occurs. This is a typical property of monostable dynamics, that fails with bistable dynamics. These lead on the contrary to what is referred to as *threshold properties*.

Next, we let the parameters *r* and *s* vary. By doing so, we observe both a monostable regime, with hair-trigger effect, and a bistable regime, with threshold properties. In order to discard from our observations rapid extinctions of bistable gene drives due to an insufficiently large initial value of *n*_*D*_, in all following simulations we use as initial value on the right (*n*_*D*_, *n*_*O*_) = (0.95, 0.05) in a wide enough interval. With such a choice of initial data, we always observe on the right the convergence of (*n*_*D*_, *n*_*O*_) to the steady state without wild-type 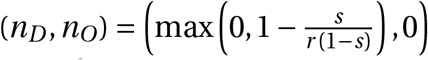.

We say that the gene drive invasion is successful, or that the gene drive is *viable*, if the invasion speed *c*_*s,r*_ is negative. On the contrary, if *c*_*s,r*_ ≥ 0, we say that the gene drive invasion fails and that the gene drive is *nonviable*: in particular, when *c*_*s,r*_ > 0, the gene drive is repelled and eliminated by the wild-type population. When introduced in the center of the spatial domain, a nonviable gene drive collapses in finite time.

In order to estimate the speed, it is convenient to have a well-defined level-set to follow. Hence we consider now the monotonicity of the traveling wave profiles. Although *N*_*D*_ is in many cases not monotonic (see again Figure 1 and Figure 2), we observe in all simulated time evolutions the following monotonicities:

**Figure 2.**
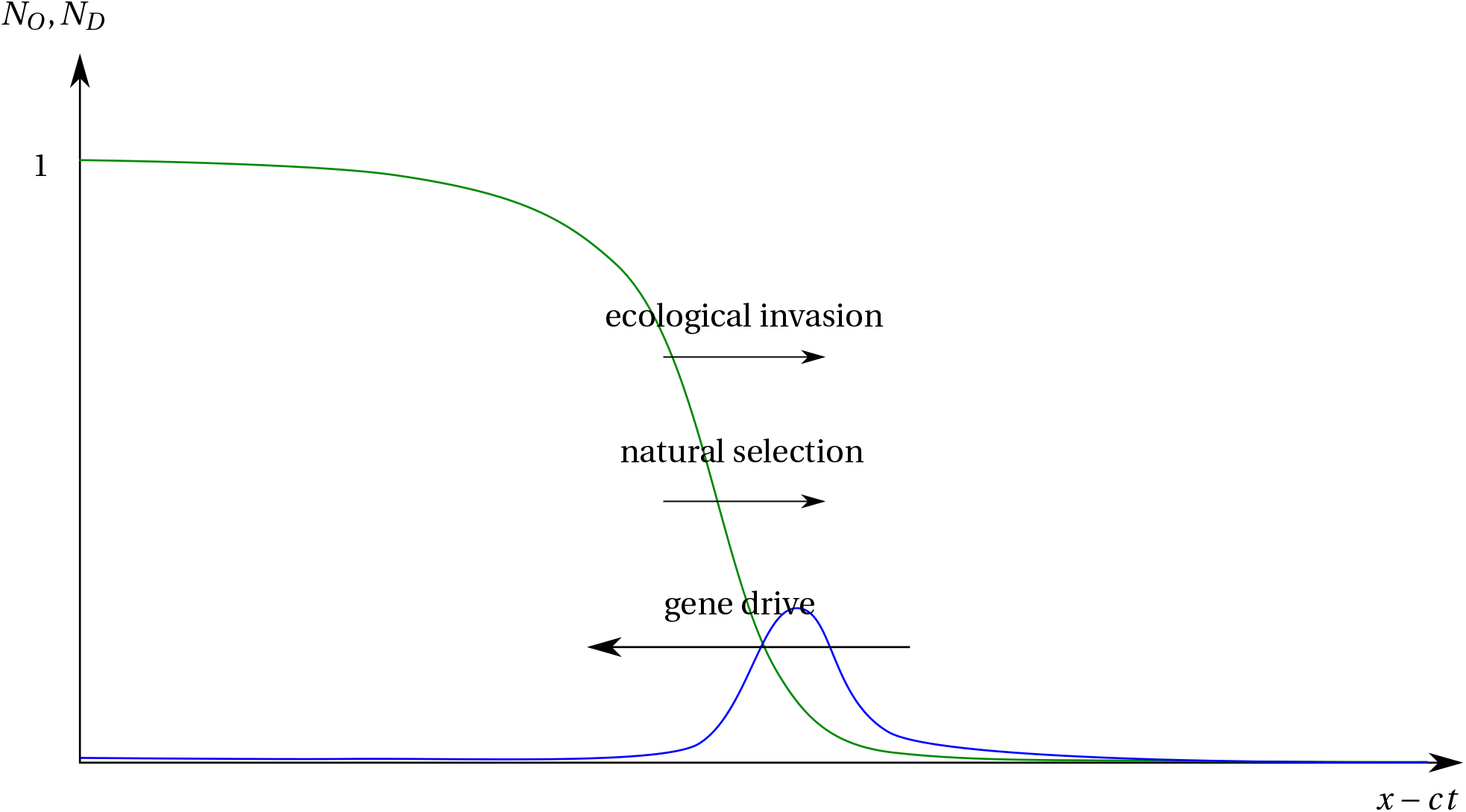
Illustration of a possible wave profile in the correct moving frame for an eradication drive. Three biological mechanisms can influence the wave speed; finding the sign of the wave speed is studying their interplay.

- the wild-type population profile *N*_*O*_ is strictly monotonic and connects 1 to 0;
- the frequency profile *P = N*_*D*_ /(*N*_*D*_ +*N*_*O*_) (plots of *p*(*t, x*) not shown here) is monotonic – but not always strictly, as we sometimes observe *P =* 0 everywhere;
- the total population profile *N = N*_*D*_+*N*_*O*_ (plots of *n*(*t, x*) not shown here) is strictly monotonic and connects 1 to 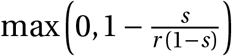.

We systematically tested the above monotonicity properties of *P* and *N* in a wide parameter range for *s* and *r*. The result is displayed on Figure 3 and confirms the preceding claim – note that 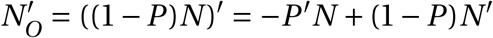 is negative as soon as −*P*′ and *N*′ are respectively nonpositive and negative, whence it suffices to check the monotonicities of *P* and *N*. Therefore we can numerically estimate the speed *c*_*s,r*_ by tracking the well-defined 1/2-level set of *n*_*O*_(*t, x*).

**Figure 3.**
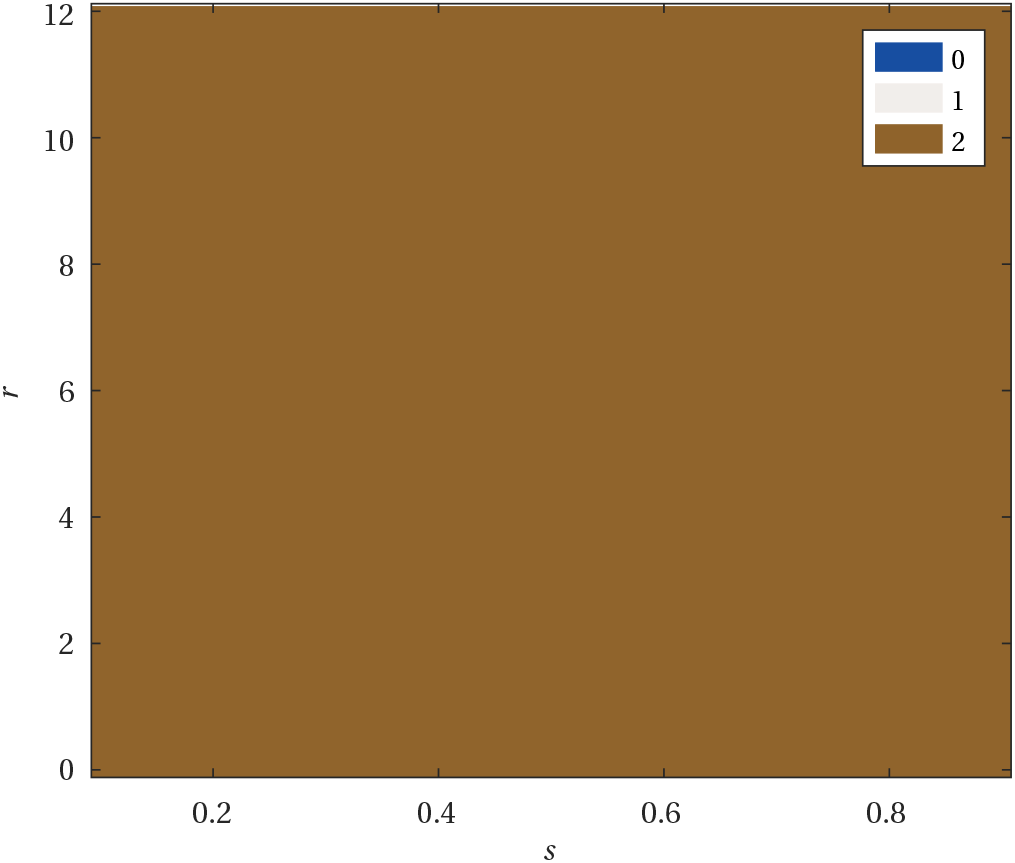
Monotonicity heatmap in the (*s, r*) plane. The output is an integer in {0, 1, 2} that approximates the number of monotonic functions in {*P, N*}. The monotonicity is tested by testing the inequalities min_*x*_ *∂*_*x*_ *p*(*T, x*)/ ∥ *∂*_*x*_ *p*(*T, x*) ∥ _*L*_∞ >− *ε* and max_*x*_ *∂*_*x*_*n*(*T, x*)/∥*∂*_*x*_*n*(*T, x*) ∥_*L*_∞ < *ε*, with *T =* 200 the final time of the simulation, when the traveling wave regime is reached. With no margin of error (*ε =* 0), the tests systematically fail due to numerical artifacts. Here, *ε =* 10^−6^ and the whole parameter range is filled in brown color: at each point (*s, r*), the normalized derivatives of *x* ↦ *p*(*T, x*) and *x* ↦ *n*(*T, x*) have both a constant sign up to a margin of error of 10^−6^.

**Figure 4.**
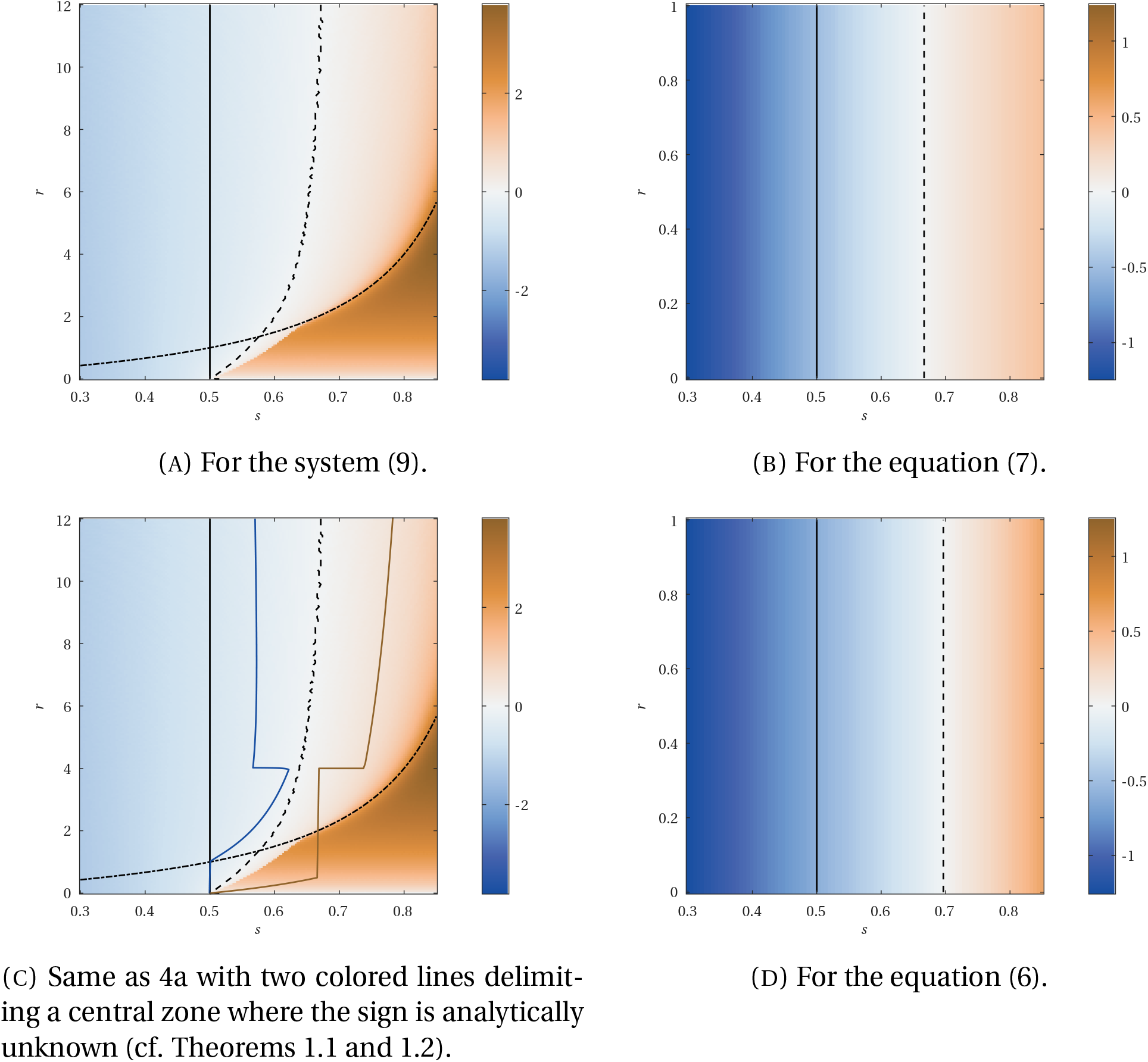
Heatmaps of *c*_*s,r*_ values in the (*s, r*) plane comparing (9) and two density-independent models (note that the scaling of the colorbar varies). Dashed curves: 0-level set. Solid black curves: monostability– bistability threshold. Dashed-dotted curve: level of *r* below which the drive is an eradication drive. As the equations (6) and (7) do not depend on *r*, the level lines on their respective figures are vertical.

Estimations of the values of *c*_*s,r*_ when *s* and *r* vary are displayed on heatmaps on Figure 4, where they are also compared to similar estimations for the density-independent equations (7) and (6) (both equations being in some way related to our model, cf. Section 1.5.1). Let us note that for the equation (7), whose reaction term is a classical cubic non-linearity, an explicit formula is known for the bistable wave speed: 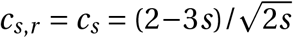. Hence its 0-level set is exactly at *s* = 2/3. For the more complicated equation (6) with a denominator, we do not know any algebraic formula for the wave speed but the 0-level set can still be computed approximately [36]: *s* ≃ 0.697. Let us also point out that despite the fact that equation (7) arises as a weak selection approximation in Section 1.5.1, we plot its values for relatively large values of *s*: in our opinion, it is interesting to compare on one hand the equation (6) and its weak selection approximation (7), as was done in Tanaka–Stone–Nelson [36], and on the other hand the equation (6) and our system (9). In particular, such a comparison clearly shows that accounting for population dynamics has a much stronger effect than accounting for strong selection.

The following comments on Figure 4a are in order.

- The wave speed *c*_*s,r*_ is (surprisingly to us) not a continuous function of (*s, r*). More precisely, a jump discontinuity divides into two parts the parameter region corresponding to nonviable eradication drives. Each part corresponds to a different kind of nonviable eradication drive: on the left-hand side of the discontinuity, we observe drives with *P* strictly increasing (case *c* > 0 of Figure 2), whereas on the right-hand side we observe drives with *P =* 0 identically. For the latter kind, the traveling wave observed numerically (see Figure 5) is actually a classical Fisher– KPP traveling wave for the *n*_*O*_ population: after having been eradicated from the right-hand side of the domain by the gene drive and having waited long enough for the eradication of the gene drive itself, the wild-type population invades the new open space at its Fisher–KPP speed 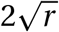. This is confirmed graphically by the fact that in the bottom right corner, the speed *c*_*s,r*_ only depends on *r*.
- Away from this discontinuity, the 0-level set of *c*_*s,r*_ seems to be an increasing, strictly convex curve, originating from (*s, r*) = (1/2, 0) and admitting a vertical straight line as asymptote. The equation of this asymptote has the form *s = s*_0_ for some undetermined *s*_0_ ∈ [0.65, 0.7]. In particular, the level set is included in {(*s, r*) | *s* ∈ [0.5, 0.7]}: if *s* < 0.5, then *c*_*s,r*_ < 0 and the gene drive invades; if *s* > 0.7, then *c*_*s,r*_ > 0 and the gene drive is nonviable. These two conclusions for the parameter range *s* ∉ [0.5, 0.7] were already valid for the density-independent equations (6) and (7) (see Figures 4b and 4d).
- There exist viable eradication drives, yet the corresponding parameter region is small, especially when intersected with the half-space defined by *s* ≥ 1/2.

**Figure 5.**
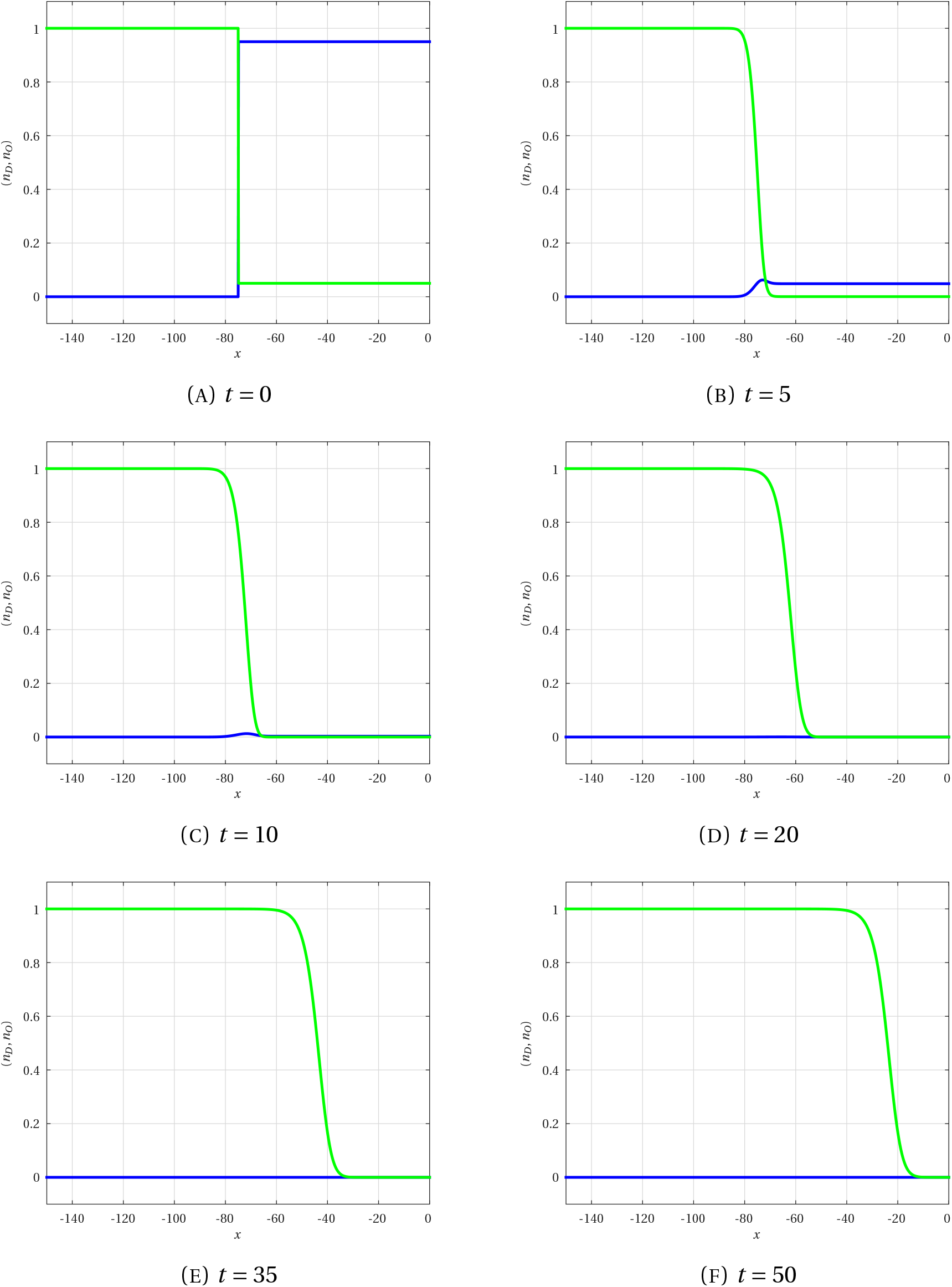
Numerical simulation of the solution of the system (9) at different times (varying time between two snapshots). Blue curve: *n*_*D*_ (*t, x*). Green curve: *n*_*O*_(*t, x*). Here *s =* 0.7, *r =* 0.5, so that by Theorem 1.1, traveling waves are trivial; moreover, (2*s* − 1)/*s =* 4/7 < 0.95/(0.95 + 0.05) = 0.95, so that the extinction of *n*_*D*_ is not due to the bistability threshold.

#### 1.6.2. Analytical results

Although finding the sign of the wave speed for a scalar bistable reaction–diffusion equation is easy (multiply the equation by the derivative of the wave profile and integrate over ℝ), it is in general a very challenging problem for bistable reaction–diffusion systems of equations. The method used in the scalar case only works for systems whose reaction term has a specific gradient form – which is not the case here – and, apart from this method, no general method is known. Systems devoid of gradient form have to be studied on a case-by-case basis and most of the time these studies lead to results on very particular cases with specific algebraic requirements on the parameters of the system. For more details on this topic, we refer to the recent review by the first author on the sign of the wave speed for two-species Lotka–Volterra competition–diffusion systems [19].

Hence we believe that, given the present state of knowledge in the mathematical analysis of reaction–diffusion systems, an explicit complete characterization of the sign of *c*_*s,r*_ is currently out of reach. We can nonetheless aim for a collection of partial results. In our opinion, the value of such results is twofold:

- on one hand, they confirm in some regions of the parameter space the numerical experiments;
- on the other hand, their proofs give precious insights on the deep structure of the equations, that might be useful in possible future work.

Regarding the existence of traveling waves (*p, n*)(*t, x*) = (*P, N*)(*x* − *ct*) with monotonic *P* and *N*, we point out first that the existence in the case *P =* 0 identically reduces to a standard question for the Fisher-KPP equation and the conclusion is well known: there exists such a t<u>r</u>aveling wave with speed *c* if and only if 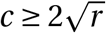 and the traveling wave with speed 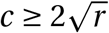 is unique (up to spatial translation). On the contrary, the case *P* ≠ 0 is a much more delicate issue here than in standard systems. On one hand, we will prove with Theorem 1.1 an explicit nonexistence result in a certain parameter range, showing that the existence for all values of (*r, s*) is simply false. On the other hand, known methods for constructing traveling waves rely either on monotonicity properties of the reaction term and super-sub-solutions or on ODE shooting arguments with stable or unstable manifolds. Yet, both methods seem inappropriate here: both systems (9) and (10) have changing monotonicities and lack 𝒞^1^ regularity as *n* → 0. We believe we might be able to obtain analytical existence results in certain parameter ranges (say, away from the eradication zone and for small values of *s* where the system has a nice KPP structure [18]), but not in the range that interests us most, namely *s* ∈ [0.5, 0.7] and in the eradication zone or close to it. Therefore we decide to leave the rigorous existence problem as an open problem^4^.

In any case, the biologically most relevant question is not the question of existence but rather the question of the direction of the propagation, namely the sign of the wave speed *c*_*s,r*_. Hence in the forthcoming Theorem 1.2 we will focus indeed on *a priori* estimates for this sign, or in other words we will study the sign of the wave speed of any existing traveling wave.

We use the variables (*p, n*) to write the following theorems; the analogous results with variables (*n*_*D*_, *n*_*O*_) can be written easily. In order to simplify the statements, we exclude traveling waves that are not monotonic or do not converge exponentially at −∞; this decision is consistent with numerical observations. We also choose to list only the results that give explicit regions of the (*s, r*) plane where the sign is known, but our proofs actually give slightly larger implicit regions.

##### Definition 1.1.

A *traveling wave solution* of (10) is a bounded nonnegative classical solution of the form (*p, n*)(*t, x*) = (*P, N*)(*x* −*ct*) satisfying:

1. *P* is nondecreasing, *N* is decreasing;
2. lim_−∞_(*P, N*) = (0, 1).

Furthermore, the traveling wave is referred to as *trivial* if *P =* 0 and *nontrivial* if *P* ≠ 0.

Obviously, a trivial, respectively nontrivial, traveling wave solution (*n*_*D*_, *n*_*O*_) of (9) is a solution such that the associated solution (*p, n*) of (10) is a trivial, respectively nontrivial, traveling wave solution itself.

Our two main results follow. Graphically, they are summarized on Figure 4c.

##### Theorem 1.1

(Nonexistence of nontrivial waves). *If* 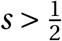 *and*

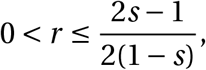

*then all traveling waves, in the sense of* *Definition 1*.*1*, *are trivial*.

*Consequently, for any traveling wave solution* (*p, n*), *n*(*t, x*) *= N* (*x* −*ct*) *is a Fisher–KPP traveling wave with speed* 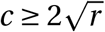.

Figure 5 is an example of spreading dynamics when 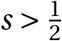 and 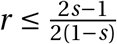.

##### Theorem 1.2

(Sign of nontrivial wave speed). *Assume* (10) *admits a nontrivial traveling wave solution* (*p, n*)(*t, x*) = (*P, N*)(*x* −*ct*), *in the sense of* *Definition 1*.*1*, *such that:*

1. *P is strictly monotonic;*
2. *P converges exponentially fast to* 0 *at* −∞.

*Then:*

1. *c* < 0 *if:*
  a. 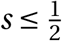; *moreover, P converges at* +*∞ to 1 and* 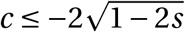;
  b. 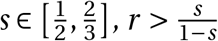, *(P,N) converges at* +*∞ to* 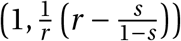 *and*

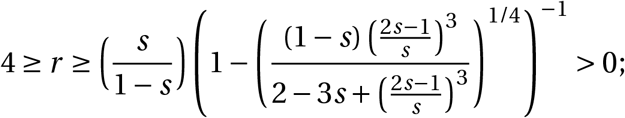
  c. 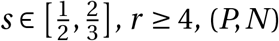 *converges at* +*∞ to* 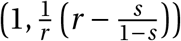 *and*

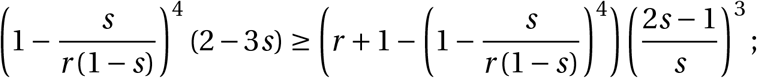
2. *c* > 0 *if:*
  a. 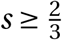 *and r*≥ 4;
  b. 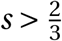 *and*

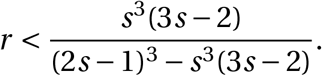

## 2. Proof of Theorems 1.1 AND 1.2

In the whole section, the spatial domain is ℝ, so that solutions of (9) or (10) are one-dimensional.

### 2.1. Proof of Theorem 1.1

#### Proposition 2.1

*Let* (*n*_*D*_, *n*_*O*_) *be a solution of* (9) *set in* (0, +∞) *×*ℝ *with n*_*D*_ (0, *x*) ≥ 0 *and n*_*O*_(0, *x*) > 0 *for all x* ∈ ℝ *and n*_*D*_ (0,•) *non-zero*.

*Then:*

1. *if* 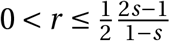, *n*_*D*_ *goes extinct spatially uniformly;*
2. *if* 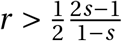 *and if n*_*D*_ (0, •) *is compactly supported, either n*_*D*_ *does not spread or it spreads at most at speed* 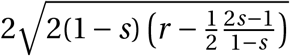.

*Proof*. Using 1 +*n*_*O*_/(*n*_*D*_ +*n*_*O*_) ≤ 2 and then 1 −*n*_*D*_ −*n*_*O*_ ≤ 1 −*n*_*D*_, we find that *n*_*D*_ satisfies

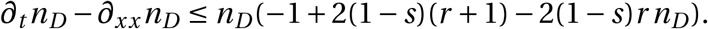

By comparison principle, 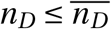, where 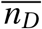 is the solution of the Fisher–KPP equation

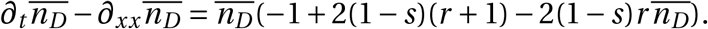

If 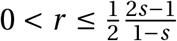, the super-solution goes extinct uniformly, and so does *n*_*D*_. Otherwise, the asymptotic speed of spreading of the super-solution, which is an upper bound for the asymptotic speed of spreading of *n*_*D*_, is exactly the speed given in the statement.□

#### Corollary 2.2.

*If* 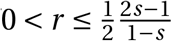, *then all traveling waves are trivial*.

*Consequently, any wave speed c satisfies* 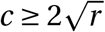.

### 2.2. Proof of Theorem 1.2

#### 2.2.1. Viable gene drives

##### Proposition 2.3.

*Assume* (9) *admits a nontrivial traveling wave solution* (*n*_*D*_, *n*_*O*_)(*t, x*) = (*N*_*D*_, *N*_*O*_)(*x* −*ct*) *satisfying*

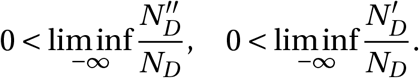

*Then, if* 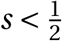, *necessarily c* < 0.

*Furthermore, if there exists λ* > 0 *such that*

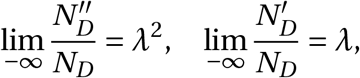

*then* 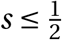 *implies* 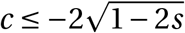.

*Proof*. Close to −∞, the wave profile *N*_*D*_ is positive and, by virtue of the assumptions, increasing 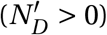 and strictly convex 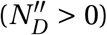. Assuming by contradiction *c* ≥ 0 and plugging these inequalities into the traveling wave equation

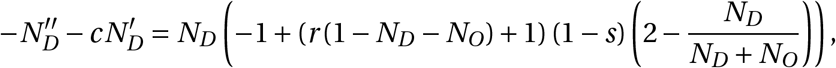

we get

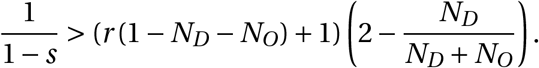

As the right-hand side converges to 2 when (*N*_*D*_, *N*_*O*_) → (0, 1), we deduce 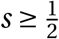.

Subsequently, assuming the exponential convergence of *N*_*D*_ with rate *λ* > 0 and passing to the limit into the equation, we discover

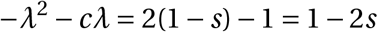

which has a positive solution if and only if *c*^2^ ≥ 4(1 − 2*s*).□

##### Proposition 2.4.

*Assume* (10) *admits a nontrivial traveling wave solution* (*p, n*)(*t, x*) = (*P, N*)(*x* −*ct*) *with increasing P*.

*Then, if* 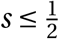, *necessarily c* < 0.

*Proof*. First, we focus on the critical case 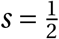.

By classification of the constant solutions of the system (10) when 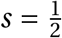, necessarily such a traveling wave admits (1, 0) or 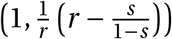 as limit at +∞ In all cases, *P* converges to 1.

We use the strict monotonicity of *N* and *P* to establish the existence of

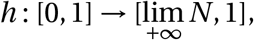

decreasing, bijective and of class *C* ^2^, such that *h*(*P*) = *N*. Using a change of variable discovered by Nadin, Strugarek and Vauchelet [26], we deduce that there exists a traveling wave *P* (*x − ct*) connecting 0 to 1 with increasing profile if and only if there exists a traveling wave solution of

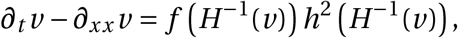

where

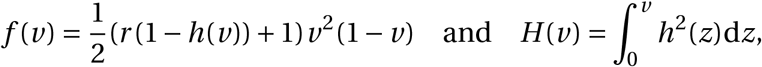

with increasing profile as well. The reaction term *g* : *v ↦ f (H*^−1^(*v)*) *h*^2^(*H*^−1^(*v)*) is non-negative and, by computing its derivatives at 0, it turns out that *g*′(0) = *f*′(0) = 0 and *g*″(0) = *f*″(0) = 1 > 0: this equation is degenerate monostable. By standard results on degenerate monostable equations with non-degenerate second derivatives (prototypical reaction term *v* ^2^(1−*v*)), there exists a traveling wave solution of speed *c* for this equation if and only if *c* ≤ *c*^⋆^, where *c*^⋆^ < 0.

Next, we consider the case 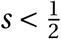. It is in fact proved similarly, except this time *g*′ (0) *= f*′ (0) > 0, so that we do not even need to look at *g*^*″*^(0).□

##### Proposition 2.5.

*Assume* (10) *admits a nontrivial traveling wave solution* (*p, n*)(*t, x*) = (*P, N*)(*x* −*ct*) *with limit* 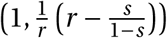 *at* +∞ *and with increasing P*.

*Then, if* 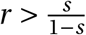 *and if one of the following conditions holds true, necessarily c* < 0:

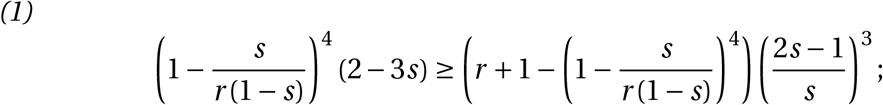

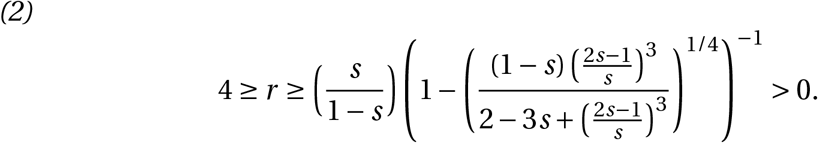

*Proof*. The case 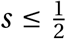 is already solved by the previous proposition, hence we assume without loss of generality 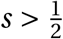. Moreover, the necessary nonnegativity of the limit implies 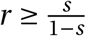; the case of equality is discarded by assumption, so that we have 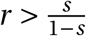.

Using again the relation *N* = *h*(*P*) and the change of variable of Nadin–Strugarek– Vauchelet [26], we find

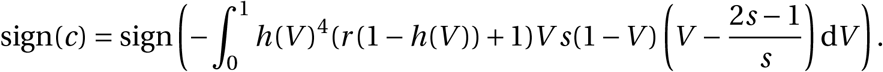

Let *I* be the integral in the right-hand side, so that sign(*c*) = − sign(*I*). Using the strict positivity of *h*, namely

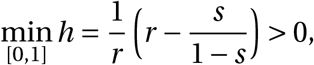

we deduce

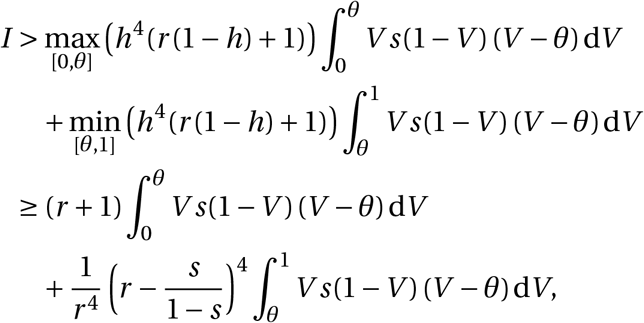

where *θ =* (2*s* − 1)/*s*. Consequently, *c* < 0 if

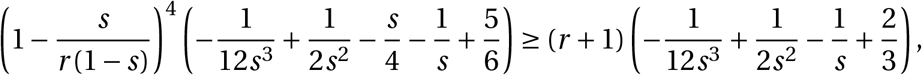

that is if

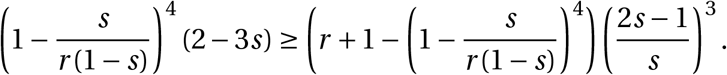

This inequality is difficult to solve explicitly, however we point out that it is both true close to 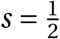 and false as *r* → +∞.

The function

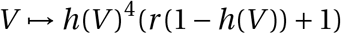

admits as derivative

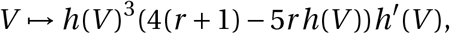

which is negative in (0, 1) if and only if *r* ≤ 4. Hence, in the particular case *r* ≤ 4, the function *h*^4^(*r* (1 −*h*) + 1) is decreasing, so that the inequality can be improved as

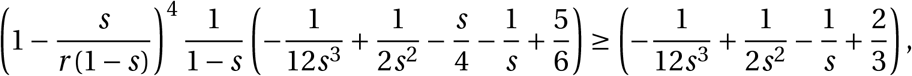

which reduces to

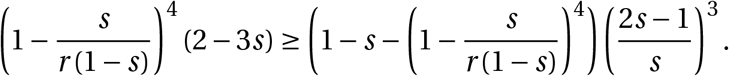

This inequality can be rewritten as

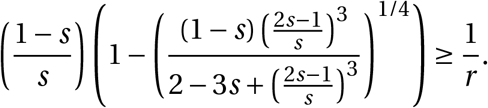

This ends the proof.□

#### 2.2.2. Nonviable gene drives

Previous propositions already confirm that nonviable gene drives can be found only in the parameter range 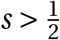.

When 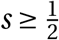, the monotonicity of (*P, N*) implies the convergence at +∞ to one constant solution among 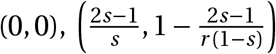 (note that this value of *n* is positive if and only if 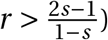 and 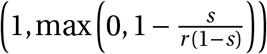.

The positivity of the speed *c* being already established in the first case (trivial traveling waves), we focus on the second and third case (nontrivial traveling waves).

##### Proposition 2.6.

*Let* (*p, n*)(*t, x*) = (*P, N*)(*x*−*ct*) *be a traveling wave solution of* (10).

*If one of the two following conditions holds true, the traveling wave is nontrivial and necessarily c* > 0:

1. *P converges at* +∞ *to* 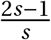 *and* 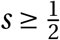;
2. *P converges at* +∞ *to* 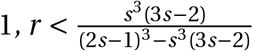 *and* 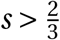;
3. *P is increasing, P converges at* +∞ *to* 1, *r* ≤ 4 *and* 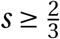.

*Proof*. Without loss of generality, assume 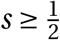.

**First condition.** The degenerate case 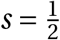 is obvious (*n*(*t,x*) is a standard Fisher–KPP traveling wave), whence we focus on 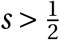.

Since *∂*_*x*_(log*n*) · *∂*_*x*_*p* = *N*′*P*′/*N≤*0, by comparison principle, 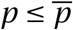 and 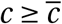, where 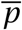 is the solution of

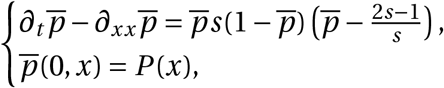

and 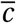 is the asymptotic speed of spreading of 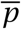. This spreading speed is larger than or equal to the associated minimal wave speed, which is positive.

**Second condition.** Since *∂*_*x*_(log*n*) · *∂*_*x*_*p* = *N*′*P*′/*N≤*0, by comparison principle, 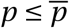 and 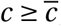, where 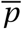 is the solution of

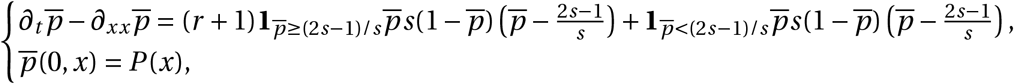

and 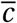 is the asymptotic speed of spreading of 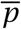.

The spreading speed of such a bistable equation is exactly its unique traveling wave speed associated with an increasing profile having limits 0 at −∞ and 1 at +∞.

Therefore the speed 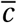 has the sign of the opposite of the integral over [0,1] of

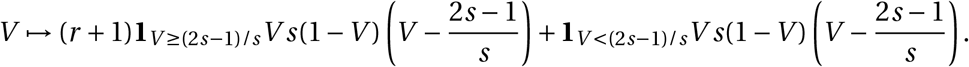

After some algebra we discover that

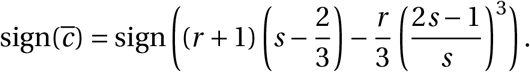

Hence 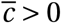 if (and only if)

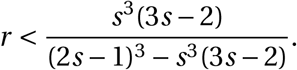

If 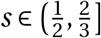, the positivity of *r* yields a contradiction, so that 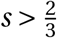 is required additionally without loss of generality. Since 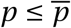 implies 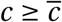, this ends the proof for the second condition.

**Third condition.** We use again the *C*^2^-diffeomorphism *h* such that *N* = *h*(*P*) and the Nadin–Strugarek–Vauchelet formula [26]:

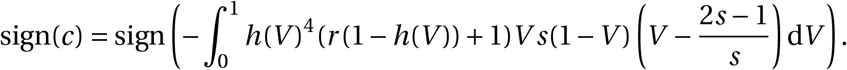

Recall that *V ↦ h*(*V*)^4^(*r* (1− *h*(*V*)) +1) is decreasing in [0, 1] if *r*≤ 4. In such a case, it can be verified that the integral *I* in the right-hand side above satisfies

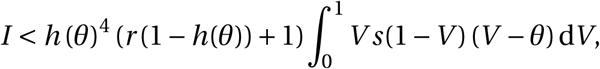

where *θ =* (2*s* − 1)/*s*. Therefore, *c* is positive if the latter integral is nonpositive, namely if *s* ≥ 2/3. □

## 3. Discussion

### 3.1. Main conclusions

In [36], Tanaka *et al*. suggested the following terminology:

- monostable gene drives, susceptible of hair-trigger effect, are *socially irresponsible gene drives*: the escape of just one individual carrying the gene drive allele from the laboratory suffices to trigger an invasion – they correspond to threshold-independent drives;
- bistable gene drives, which can never invade if released accidentally in small quantities, are on the contrary *socially responsible gene drives* – they correspond to high-threshold drives.

This terminology can be combined with our viable/nonviable terminology: all socially irresponsible gene drives are viable but socially responsible gene drives can, *stricto sensu*, be viable or nonviable. Of course, practical gene drives should all be both viable and socially responsible. In Tanaka–Stone–Nelson’s density-independent model [36], this meant simply 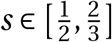. However, in the model presented here, neither social responsibility (the drive is a threshold-dependent drive) nor viability (the drive is able to invade a wild-type population) can be easily and explicitly characterized with analytic methods. Numerical simulations become essential in understanding the interplay between the two parameters *s* and *r*.

Our model also shows that a gene drive can achieve complete eradication only if *r* is small enough. This leads to a particulary striking conclusion: only very specific choices of *s* and *r*, corresponding to a small compact region in the (*s, r*)-plane, can lead to socially responsible and viable eradication gene drives.

More generally, our model shows that the invasion of an eradication drive but also of any gene drive achieving only partial population suppression can be slowed down, stopped or even reversed by the opposing demographic advection term. Thus population dynamics matters for any gene drive affecting population size, even slightly, and should be taken into account. We however find that threshold-independent drives can spread spatially even when they lead to population suppression (thereby extending a result found by Beaghton *et al*. [5] for driving-Y to homing-based gene drives): their monostability property remains true, their hair-trigger effect is still observed numerically and the demographic advection term only slows them down but never stops them. For practical applications, this feature is especially dangerous if one thinks of threshold-independent eradication drives. These should definitely be approached with the highest caution.

Thus, ignoring population dynamics and only focusing on allele frequencies for spatial models of gene drive [36] should be limited to replacement drives, that correspond for instance to the weak selection framework (small values of *s*) or to very fast population dynamics (large values of *r*). We refer to the discussion in Section 1.5.1 on the relation between our model and the two density-independent equations (6) and (7).

### 3.2. Mathematical open problems worthy of attention

Characterizing analytically the asymptotics *r* → 0 and *r* → +∞, namely:

- the transition at *s =* 1/2 when *r ≃* 0, from nonexistent traveling waves to existent monostable traveling waves, without intermediate bistable regime;
- the vertical asymptote of the level line {*c*_*s,r*_ = 0}, its precise value in [0.65, 0.7] and its relation with Strugarek–Vauchelet [35];

are two natural but difficult questions, that are left for further studies.

The main open problem with the systems (9) and (10) is the existence of nontrivial traveling waves. We briefly explained in Section 1.6.2 why this problem is very difficult: on one hand, the nonexistence result of Theorem 1.1 means that there will necessarily be requirements on the parameters *s* and *r*, and on the other hand known methods for proofs of existence are likely to be inappropriate, especially in the most interesting parameter range *s* ∈ [1/2, 2/3].

Some methods to construct traveling waves do not guarantee the monotonicity of the constructed profiles. Numerically, we only observed monotonic profiles *N* and *P*, but we did not manage to prove the *a priori* monotonicity of these profiles. Thus at this point it remains possible that traveling waves with non-monotonic profiles coexist with the observed monotonic ones.

These questions of existence lead to another related crucial issue: the stability properties of traveling waves^5^. Indeed, we know that trivial traveling waves always exist. When a trivial and a nontrivial wave coexist, can we predict theoretically which one is selected? The numerical experiments are ambivalent here. On one hand, they lead naturally to the following (rough) conjecture:

*Conjecture* 3.1. In the whole parameter region where monotonic nontrivial traveling waves are numerically selected,

1. trivial traveling waves are unstable;
2. monotonic nontrivial traveling waves exist and are stable.

But on the other hand, it is numerically very challenging to distinguish cases where only trivial waves exist from cases where the two kinds coexist but the trivial ones are the only stable ones. Hence, at this point, it remains unclear whether the whole bottom right region of Figure 4a corresponds to a nonexistence result or not. We strongly believe that Theorem 1.1 is not optimal, but we do not know if we can expect the optimal result to cover the whole region where 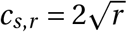.

Finally, it would of course be interesting to improve our theoretical knowledge on the sign of the speed of nontrivial traveling waves. A direction that might be fruitful is the following: find upper or lower estimates for the decreasing function *h* : [0, 1] → [0, 1] defined by *h*(*P*) *= N* and plug these estimates into the Nadin–Strugarek–Vauchelet formula [26] that we used repeatedly in the proofs:

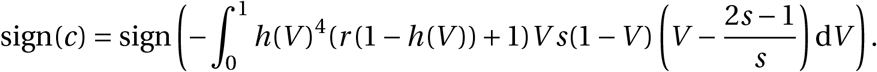

Let us point out that a second similar formula can be obtained by integration by parts of the equation on *N* multiplied by *N*′:

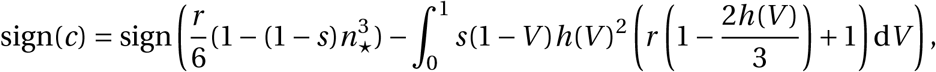

where 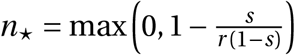. Unfortunately we did not manage to deduce anything interesting from this formula. But, again, it might prove fruitful once estimates on *h* are established.

### 3.3. Effect of stochasticity

We checked the robustness of our results by running stochastic simulations of the model given in system (9). More precisely, our stochastic model is a variant of our deterministic model implemented as a modified Gillespie algorithm, with Brownian motion of individuals, numbers of offspring drawn according to a Poisson law and death times drawn according to an exponential law. Each simulation is stopped when one allele completely disappears from the spatial domain or when a given final time is reached (even if no allele has disappeared yet). Space was discretized into 100 demes connected by dispersal to the nearest neighboring sites, with emigration probability equal to 0.1; the carrying capacity of a deme is *K =* 1000. The simulation codes are currently available online at https://plmlab.math.cnrs.fr/fdebarre/2021_DriveInSpace.

These simulations confirmed the results of the deterministic version of the model (compare Figure 6 to Figure 4a).

**Figure 6.**
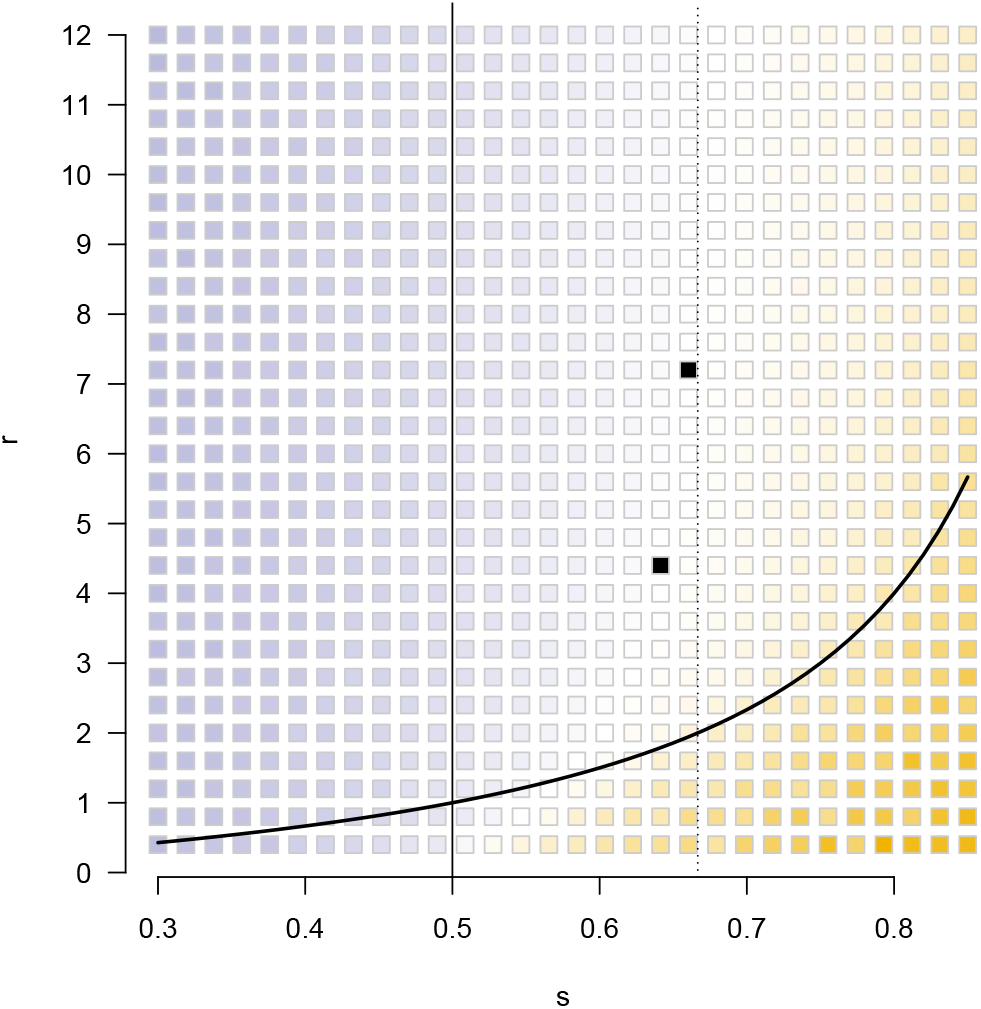
Heatmap in the (*s, r*) plane for the stochastic simulations. Each point is the color-coded outcome of one stochastic simulation run with the corresponding (*s, r*) parameters. Blue: the simulation is stopped by the extinction of *O*, yellow: the simulation is stopped by the extinction of *D*, black: no extinction occurs before the final time. The more intense the color, the faster the extinction of the last individual carrying the first lost allele. The thick curve represents the eradication threshold.

### 3.4. Genericity of this “wave reversal by opposing demographic advection” phenomenon

#### 3.4.1. Other choices of birth and death rates

Numerically, several other cases were considered for *B* and *N*

1. Constant death rate, weak or strong Allee effect on the wild-type dynamics with threshold *a* < 1: *D*(*n*) = 1, *B* (*n*) = max(*r* (1 −*n*)(*n* − *a*) + 1, 0) (as a birth rate, *B* has to be nonnegative). The Allee effect is weak if *a* ∈ (−1, 0] and strong if *a* ∈ (0, 1) (if *a* ≤ −1, there is again no Allee effect). The eradication condition is 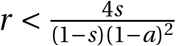.
2. Constant birth rate, logistic wild-type growth: *B* (*n*) *= r* + 1, *D*(*n*) = 1 + *r n*. Again, the eradication condition is 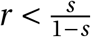.
3. Constant birth rate, weak or strong Allee effect with threshold *a* ∈ (−1, 1) on the wild-type growth: *B* (*n*) *= r* + 1, *D*(*n*) = 1 + *r* + *r* (*n* − 1)(*n* − *a*). The eradication conditions are (1 − *a*)^2^ ≤ 4*s* or 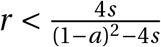. Note that the first condition does not depend on *r*.

The resulting heatmaps can be observed on Figure 7. Interestingly, many features of Figure 4a remain true: the 0-level set is an increasing graph, it is stuck in the parameter range defined by the two thresholds *s =* 1/2 and *s =* 2/3 of the density-independent equation (7). However, the strong Allee effect on *B* seems to break the strict convexity of the 0-level set and the strong Allee effect on *D* seems to simply the picture in the sense that all nonviable gene drives satisfy *P =* 0.

**Figure 7.**
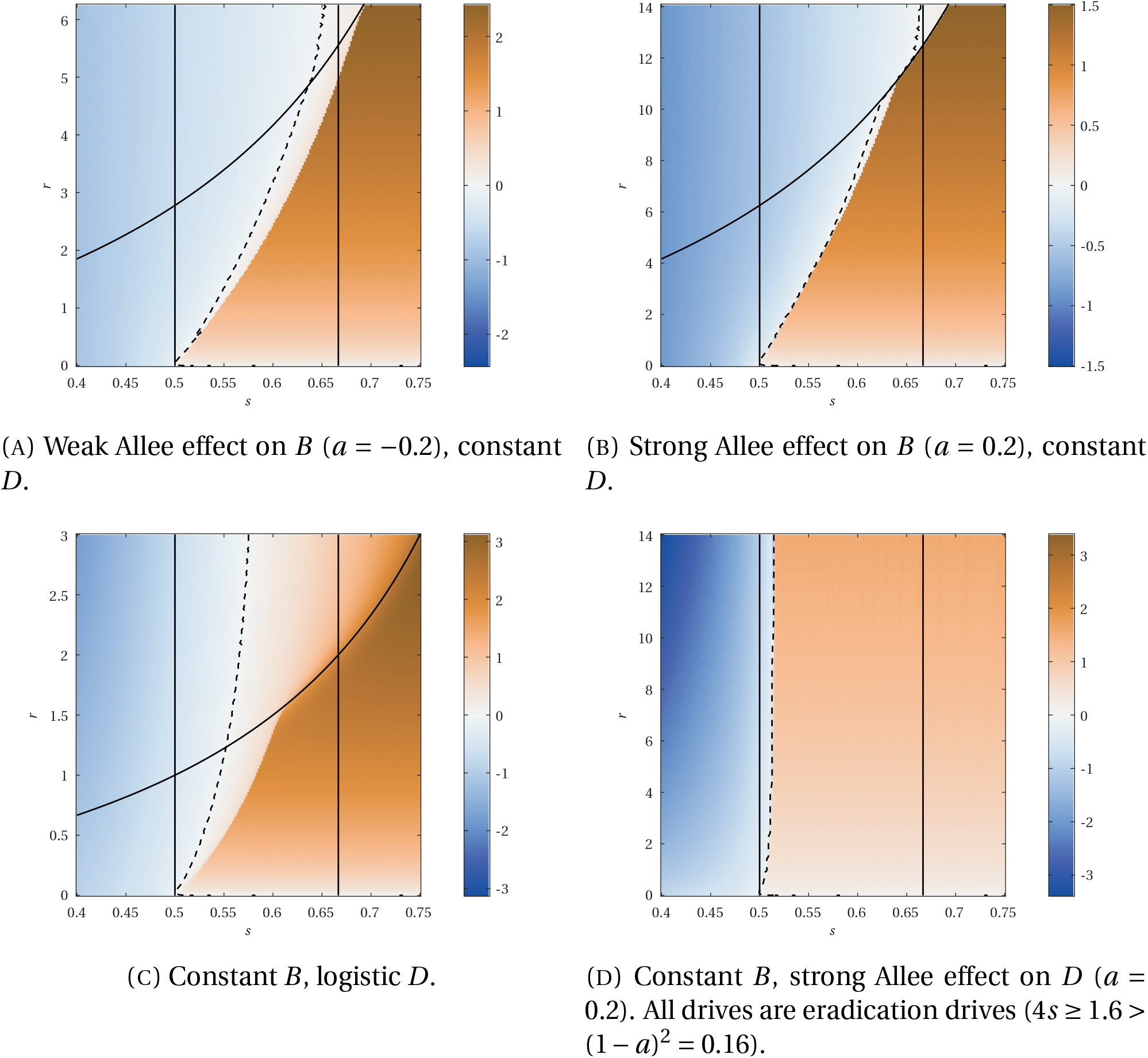
Heatmaps of *c*_*s,r*_ values in the (*s, r*) plane for different choices of *B* and *D* in (2). Dashed curves: 0-level set. Strictly convex curves: eradication threshold. The associated density-independent dynamics are given by (7), with thresholds *s =* 1/2 and *s =* 2/3 (see Figure 4b).

We point out that many analytical techniques used in the proofs of Theorems 1.1 and 1.2 can be successfully applied to those cases. For the sake of brevity, we do not give details.

#### 3.4.2. Other gene drive models

The same numerical method can be applied to different gene drive models: different particular cases of the system (1), particular cases of the analogous model obtained when assuming that the gene conversion (homing) occurs in the germline [28, 31], or entirely different models [11, 36, 38, 39]. We illustrate this possibility with two examples on Figure 8. Again, the main features of Figure 4a are preserved. However we point out that our analytical tools strongly rely upon the fact that the system is a two-component system: most of them would be unapplicable in a three-component setting. This implies that models where three genotypes *OO, OD* and *DD* have to be tracked simultaneously cannot be studied this way. Our analytical approach seems to be limited to gene drives with perfect conversion and conversion in the zygote, for which heterozygous individuals can be safely ignored, or to models reduced by means of some approximation (e.g. Hardy-Weinberg proportions as in discrete-time models).

**Figure 8.**
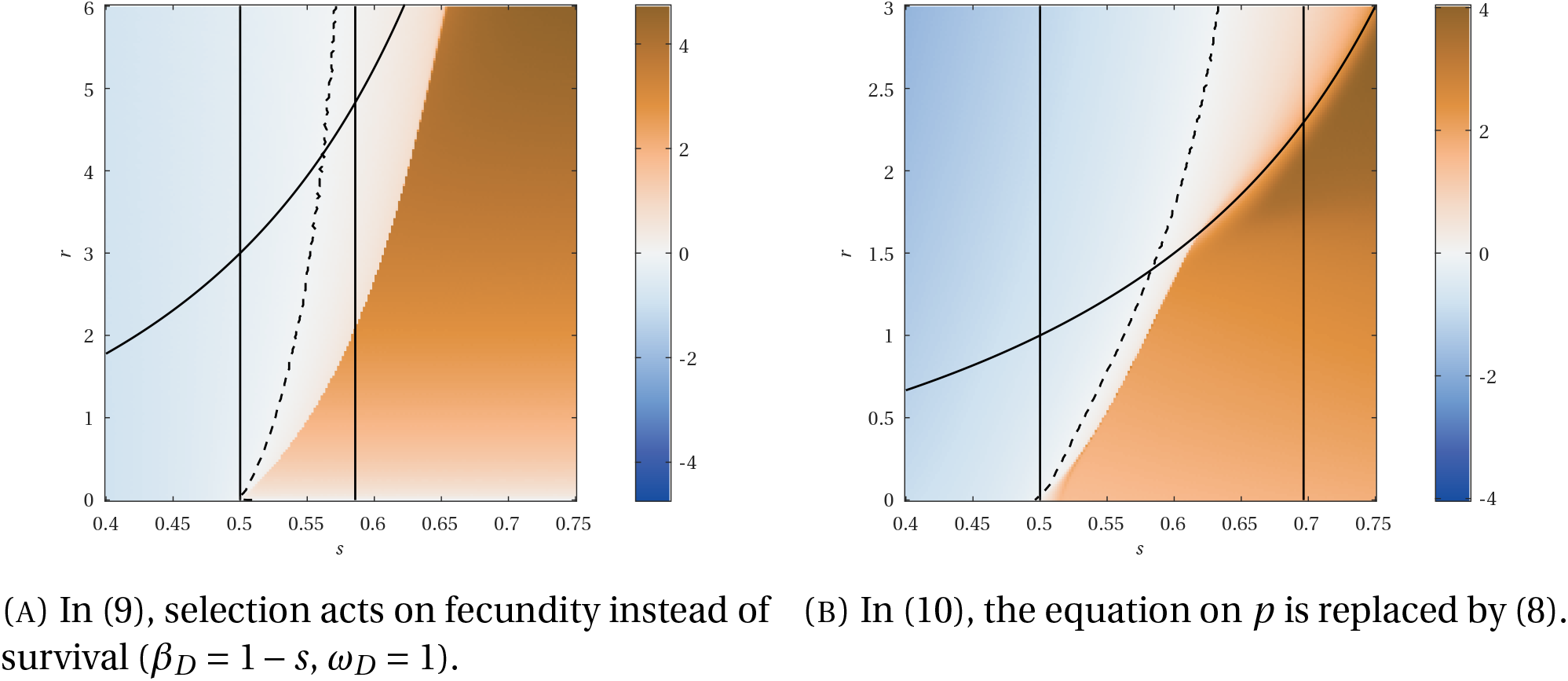
Heatmaps of *c*_*s,r*_ values in the (*s, r*) plane for different gene drive models. Dashed curves: 0-level set. Strictly convex curves: eradication threshold. Left vertical line: monostability–bistability threshold for the density-independent dynamics. Right vertical line: 0-level set of the speed of the density-independent dynamics.

#### 3.4.3. Extension to a bistable mosquito–Wolbachia model

Finally, the same ideas can also be applied to different models from mathematical biology that share the common feature of leading to a two-component bistable reaction–diffusion system where one population with low carrying capacity tries to invade another population with high carrying capacity. Indeed, such a model will again involve an opposing advection term of the form −2∇(log *n)* · ∇*p* and it should be clear at this point that this term is the main responsible for the qualitative features of Figure 4a.

As an example, we consider a Wolbachia infection in a mosquito population, as described in [4, 26, 34]: the infection is transmitted vertically, perfectly, through mothers. The infection reduces fertility of mothers by a factor *f*_*w*_. Hatching rate is reduced by a factor *ω*_*H*_ for crosses involving an infected father and an uninfected mother. All other crossings have the same hatching rates. Assuming equal sex ratios, no sex differences in death rate and no differences in diffusion rate, we have

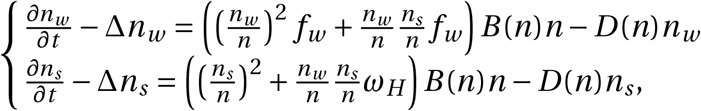

where *n*_*w*_ and *n*_*s*_ are the densities of infected and uninfected individuals, respectively, and *n = n*_*w*_ + *n*_*s*_ is the total population density. Rewriting this system in terms of *n* and 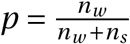, we obtain

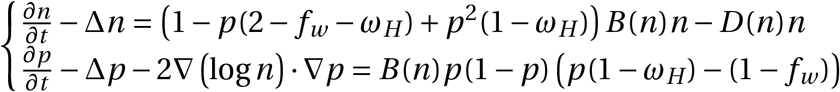

The interior equilibrium for *p*,

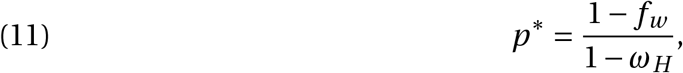

is admissible (*i*.*e*., between 0 and 1) when *ω*_*H*_ ≤ *f*_*w*_ ; it is unstable. With this example, we can indeed have a wave towards lower densities (a fully infected population is smaller than a fully uninfected population).

Numerical simulations for this model are displayed on Figure 9. Again, this figure shares many similarities with Figure 4a. This illustrates once more that the important term in this whole class of models when concerned about spreading properties is the opposing advection term.

**Figure 9.**
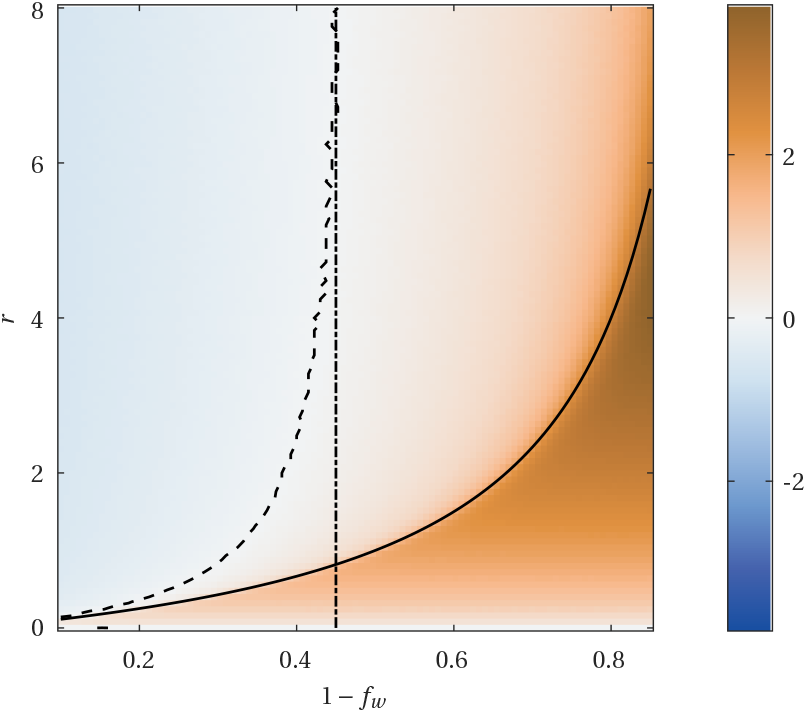
Heatmaps of 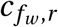 values in the (1 − *f*_*w*_, *r*) plane for the Wolbachia system with *ω*_*H*_ = 0.1. Dashed curve: 0-level set. Solid curve: level of *r* below which Wolbachia leads to extinction. Dashed-dotted line: 0-level set of the speed of the density-independent dynamics.

### 3.5. Biological implications

We found that a threshold-independent homing drive would spread spatially, even if it leads to the eradication of the target population. This extends the results of Beaghton *et al*. [5], obtained by a driving-Y, to another type of gene drive. For threshold-dependent drives, however, we find that the parameter space allowing for spatial spread is more restricted than what is found in models neglecting changes in population size and the resulting opposing demographic advection of individuals [36]. These simpler models find that a drive spreads as long as the release threshold is sufficiently low (lower than 1/2 in [3, 4], lower than a value close to 0.565 in [36]) and the initial condition is sufficiently large. In contrast, we find that these conclusions remain true for pure replacement drives but that spatial spread is limited for suppression or eradication drives, all the more as suppression is strong. These conclusions can be extended to other, non-drive, systems for population control such as Wolbachia.

The parameter range displayed on Figure 4 and other figures for the Malthusian growth rate, namely *r* ∈ [0, 12], seems to be biologically realistic and relevant. Indeed, we found *r =* 11.5 in [41] for *Aedes aegypti*, 2 ≤ *r* ≤ 9.8 in [13] for *Aedes aegypti, r =* 1.4 in [25] for *Anopheles gambiae, r =* 0.2 in [22] for *Anopheles stephensi* ^6^, *r =* 1.3 in [32] for *Aedes albopictus*.

We have hence characterised a new reason for the failure of spatial spread of suppression drives, in the form of opposing demographic advection. This phenomenon was expected given previous work on spatial dynamics of alleles (as reviewed in [12]), but we clarify conditions under which it occurs. Other models of spatial spread, and in particular individual-based models, had already identified some reasons why the spatial spread of a suppression drive may fail. If the drive suppresses the local population too much and if the density target population is spatially heterogeneous, the drive may go extinct locally with the eradication of a local subpopulation before it could spread to other locations [29]. Strategies relying on the eradication of the target population are also limited by the potential recolonization of emptied locations by wild-type individuals [9,29,30] (such recolonizations can also be observed in our stochastic simulations). Finally, the evolution of resistance to the drive itself, which already hinders the success of gene drives in well-mixed populations [37], also affects their spatial spread [6].

Our model was derived under limiting assumptions, including a 100% homing rate, and either homing taking place very early in development or the drive being dominant. Gene drives currently being designed in laboratories do not exactly match these assumptions. While we are pessimistic that analytical results can be obtained when these assumptions are relaxed, future numerical or computational (individual-based) studies will be useful to assess the generality of our findings. The results of individual-based simulations of the spatial spread of underdominance gene drive systems [10] are encouraging. The authors indeed found limited spatial spread even in the case of *p*\* < 1/2, in simulations initiated with the left half of the arena occupied by drive homozygotes and the right half by wild-type homozygotes (2L2T system; Fig. 3F) – initial conditions similar to ours.

To conclude, this work highlights the importance of not neglecting demographic dynamics in evolutionary models when the spread of an allele affect the size of a population.

## Acknowledgements

We are grateful to the INRAE MIGALE bioinformatics facility (MIGALE, INRAE, 2020. Migale bioinformatics Facility, doi: 10.15454/1.5572390655343293E12) for providing computing resources. FD is funded by an Agence Nationale de la Recherche JCJC grant TheoGeneDrive ANR-19-CE45-0009-01.

## Footnotes

1 For any *K* > 0, the pair (*n*_*D*_, *n*_*O*_) with birth and death rates *B* and *D* satisfies the same system as the pair 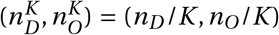 with birth and death rates *B*^*K*^ (*n*) *= B* (*Kn*), *D*^*K*^ (*n*) *= D*(*Kn*). Hence the normalization can be done without loss of generality and all forthcoming results do not depend on the wild-type carrying capacity.

2 The shared history between reaction–diffusion PDEs and population genetics is ancient: we remind the reader that both Fisher [17] and Kolmogorov, Petrovsky and Piskunov [23] introduced the equation that is now famously called the Fisher–KPP equation as a population genetics model.

3 It was precisely the aim of Strugarek and Vauchelet [35] to make such formal statements rigorous.

4 The empirical proof of existence provided by our numerical experiments might be sufficiently convincing for many non-mathematicians.

5 This whole paragraph is intentionally vague regarding the notion of stability, as we do not want to enter into details here. Of course, an unambiguous clarification would be necessary before any attempt at a rigorous proof.

6 From [13, 22, 25, 41], *r* is deduced by using the formula *r = b*/*d* − 1, where *b* is the constant adult reproduction rate (namely, the birth rate corrected by taking into account juvenile mortality), *d* is the constant adult death rate, and *b* and *d* are given in the same unit, say day^−1^. Compared with the classical formula *r = b* −*d*, here we divide by *d* to account for the fact that in system (9) the time unit is modified so that the death rate is scaled to 1.

## Notes

https://plmlab.math.cnrs.fr/fdebarre/2021_DriveInSpace

### Competing Interest Statement

The authors have declared no competing interest.

